# Conditional Gene Expression in *Magnaporthe oryzae* via Fungal Nitrate Reductase Promoter Replacement

**DOI:** 10.1101/2025.10.30.685542

**Authors:** Justin King, Mostafa Rahnama

## Abstract

Functional genomic studies rely on controlled gene expression, which is challenging for essential genes or those active only under specific conditions. We provide a detailed protocol for generating *Magnaporthe oryzae* (synonym of *Pyricularia oryzae*) with conditional *BUF1* gene expression. This method uses targeted promoter replacement to substitute the native *BUF1* promoter with the nitrogen-responsive promoter. This approach enables nitrogen source-dependent control of expression in *M. oryzae*, leading to observable pigmentation changes that facilitate easy phenotypic screening.

## BEFORE YOU BEGIN

### CRITICAL

To prevent contamination, all procedures involving fungal spores and cultures should be carried out in a sterile environment, such as a biosafety cabinet, using sterilized materials and equipment.

### Innovation

Functional genomic studies of essential genes in haploid fungi require conditional gene expression systems, as traditional gene deletion approaches often yield non-viable mutants. While conditional promoter replacement (CPR) strategies using nitrate reductase promoters have been established in other fungi ^1, 2^, comprehensive protocols for implementing these systems in *Magnaporthe oryzae* (synonym of *Pyricularia oryzae*), a critical model for plant-pathogen interactions and rice blast disease—have been lacking. In this study, we present a detailed and reproducible methodology for targeted promoter replacement in the *M. oryzae*, validating the *pMoNIA1* promoter as a functional tool for nitrogen-responsive conditional gene expression in its native fungal host. We provide a complete workflow for Agrobacterium-mediated transformation and homologous recombination-based promoter replacement ^3^, including optimized media formulations, transformation conditions, and comprehensive troubleshooting strategies that maximize efficiency and reproducibility across experiments. To demonstrate the system’s functionality, we strategically employ *BUF1*, a melanin biosynthesis gene whose down-regulation produces a distinct and easily observable color change from gray/black to buff/tan ^4^, providing clear visual confirmation that *pMoNIA1*-based CPR operates as intended in *M. oryzae*. This validated system enables reversible, media-controlled gene expression with tight regulation. By demonstrating successful promoter replacement, nitrogen-dependent regulation, and predictable phenotypic consequences with *BUF1*, we establish a robust and accessible tool for the *M. oryzae* research community to perform functional analysis of essential genes, genes with pleiotropic effects, or those required at specific developmental stages—genes that would otherwise be intractable through conventional genetic approaches.

### Institutional permissions

All experiments involving *M. oryzae* must be performed in accordance with relevant institutional and national guidelines and regulations for handling plant pathogenic fungi. Researchers must acquire permissions from their relevant institutions prior to commencing experiments.

### Fungal Culture and Maintenance

1. Maintain *M. oryzae* strain on Oatmeal Agar (OM) plates at 25°C in a growth chamber under light.
  - **Note:** Regular subculturing (every 2-3 weeks) is recommended to maintain vigorous growth and sporulation.
2. Prepare fresh fungal cultures for spore suspension preparation.

## KEY RESOURCES TABLE

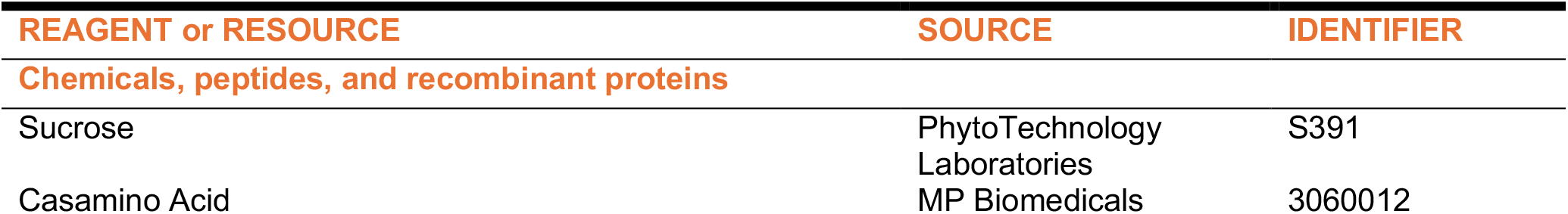
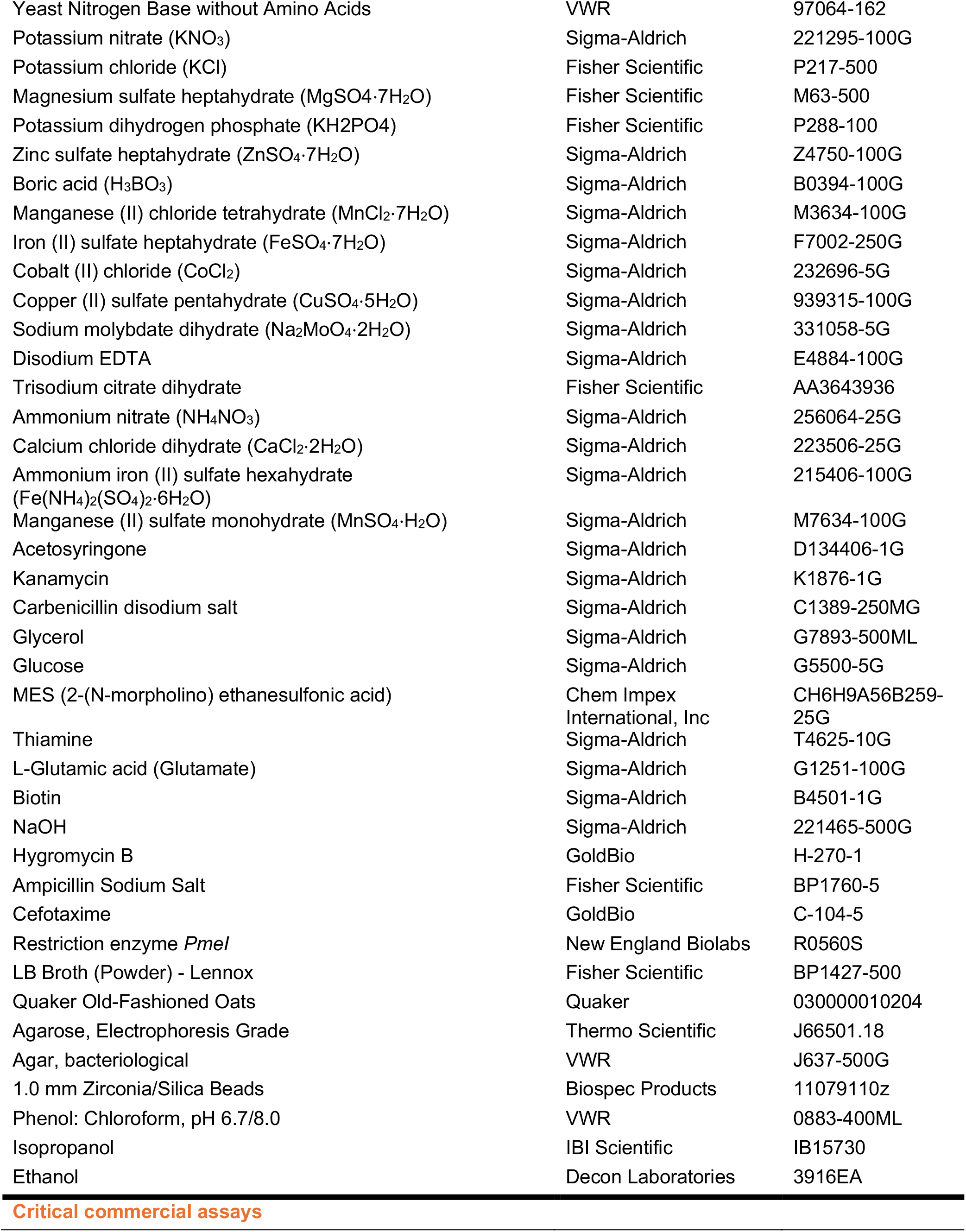
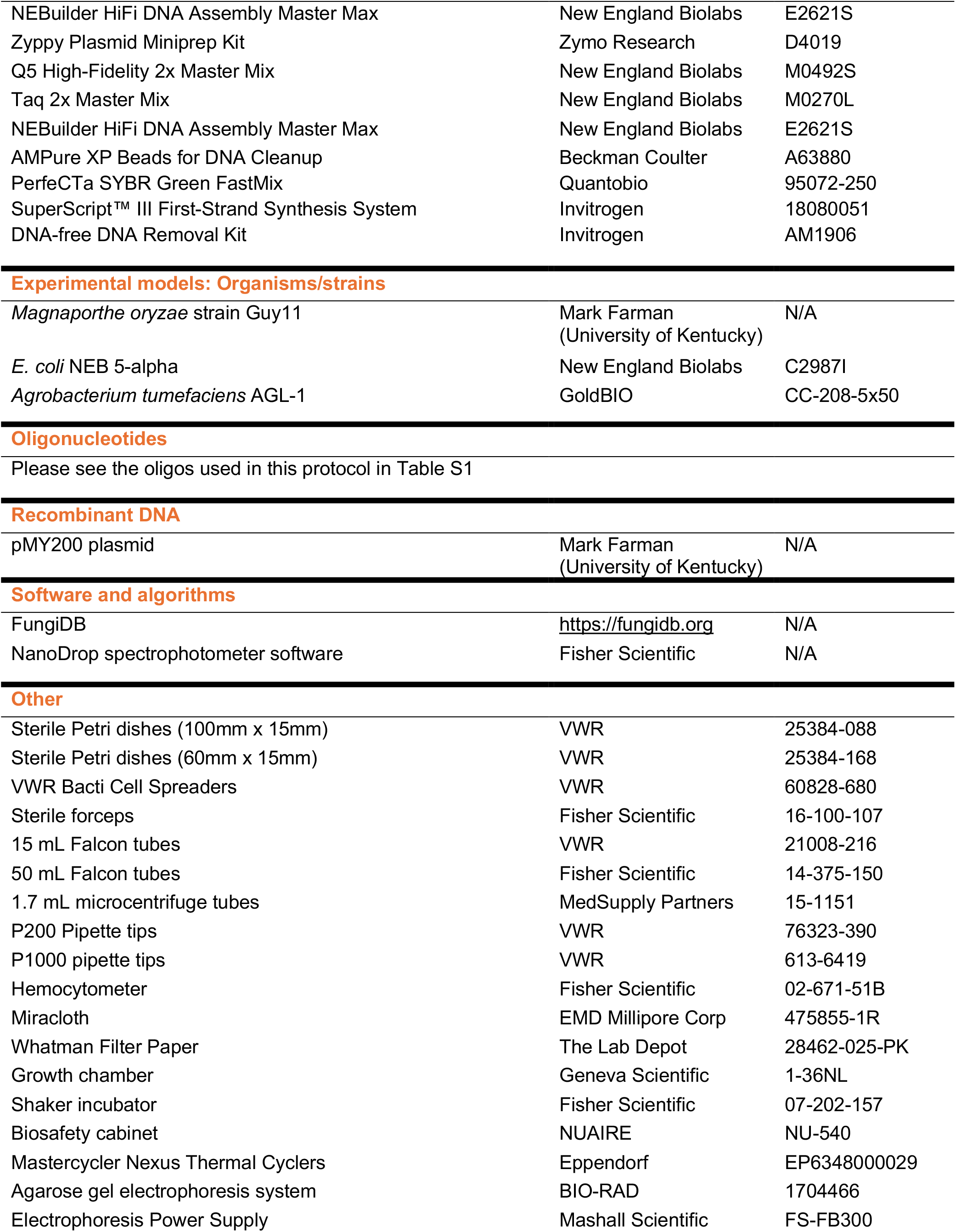
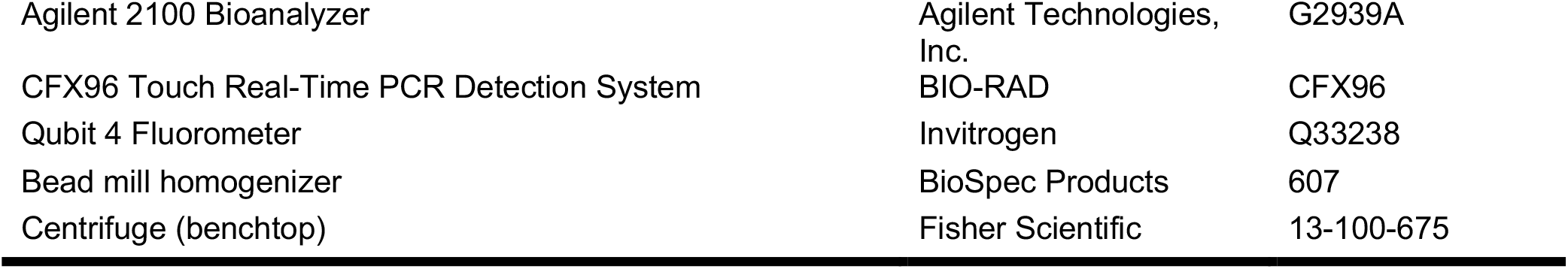

| REAGENT or RESOURCE | SOURCE | IDENTIFIER |
| --- | --- | --- |
| **Chemicals, peptides, and recombinant proteins** |  |  |
| Sucrose | PhytoTechnology Laboratories | S391 |
| Casamino Acid | MP Biomedicals | 3060012 |
| Yeast Nitrogen Base without Amino Acids | VWR | 97064-162 |
| Potassium nitrate (KNO<sub>3</sub>) | Sigma-Aldrich | 221295-100G |
| Potassium chloride (KCl) | Fisher Scientific | P217-500 |
| Magnesium sulfate heptahydrate (MgSO<sub>4</sub>·7H<sub>2</sub>O) | Fisher Scientific | M63-500 |
| Potassium dihydrogen phosphate (KH<sub>2</sub>PO<sub>4</sub>) | Fisher Scientific | P288-100 |
| Zinc sulfate heptahydrate (ZnSO<sub>4</sub>·7H<sub>2</sub>O) | Sigma-Aldrich | Z4750-100G |
| Boric acid (H<sub>3</sub>BO<sub>3</sub>) | Sigma-Aldrich | B0394-100G |
| Manganese (II) chloride tetrahydrate (MnCl<sub>2</sub>·7H<sub>2</sub>O) | Sigma-Aldrich | M3634-100G |
| Iron (II) sulfate heptahydrate (FeSO<sub>4</sub>·7H<sub>2</sub>O) | Sigma-Aldrich | F7002-250G |
| Cobalt (II) chloride (CoCl<sub>2</sub>) | Sigma-Aldrich | 232696-5G |
| Copper (II) sulfate pentahydrate (CuSO<sub>4</sub>·5H<sub>2</sub>O) | Sigma-Aldrich | 939315-100G |
| Sodium molybdate dihydrate (Na<sub>2</sub>MoO<sub>4</sub>·2H<sub>2</sub>O) | Sigma-Aldrich | 331058-5G |
| Disodium EDTA | Sigma-Aldrich | E4884-100G |
| Trisodium citrate dihydrate | Fisher Scientific | AA3643936 |
| Ammonium nitrate (NH<sub>4</sub>NO<sub>3</sub>) | Sigma-Aldrich | 256064-25G |
| Calcium chloride dihydrate (CaCl<sub>2</sub>·2H<sub>2</sub>O) | Sigma-Aldrich | 223506-25G |
| Ammonium iron (II) sulfate hexahydrate (Fe(NH<sub>4</sub>)<sub>2</sub>(SO<sub>4</sub>)<sub>2</sub>·6H<sub>2</sub>O) | Sigma-Aldrich | 215406-100G |
| Manganese (II) sulfate monohydrate (MnSO<sub>4</sub>·H<sub>2</sub>O) | Sigma-Aldrich | M7634-100G |
| Acetosyringone | Sigma-Aldrich | D134406-1G |
| Kanamycin | Sigma-Aldrich | K1876-1G |
| Carbenicillin disodium salt | Sigma-Aldrich | C1389-250MG |
| Glycerol | Sigma-Aldrich | G7893-500ML |
| Glucose | Sigma-Aldrich | G5500-5G |
| MES (2-(N-morpholino) ethanesulfonic acid) | Chem Impex International, Inc | CH6H9A56B259-25G |
| Thiamine | Sigma-Aldrich | T4625-10G |
| L-Glutamic acid (Glutamate) | Sigma-Aldrich | G1251-100G |
| Biotin | Sigma-Aldrich | B4501-1G |
| NaOH | Sigma-Aldrich | 221465-500G |
| Hygromycin B | GoldBio | H-270-1 |
| Ampicillin Sodium Salt | Fisher Scientific | BP1760-5 |
| Cefotaxime | GoldBio | C-104-5 |
| Restriction enzyme *PmeI* | New England Biolabs | R0560S |
| LB Broth (Powder) - Lennox | Fisher Scientific | BP1427-500 |
| Quaker Old-Fashioned Oats | Quaker | 030000010204 |
| Agarose, Electrophoresis Grade | Thermo Scientific | J66501.18 |
| Agar, bacteriological | VWR | J637-500G |
| 1.0 mm Zirconia/Silica Beads | Biospec Products | 11079110z |
| Phenol: Chloroform, pH 6.7/8.0 | VWR | 0883-400ML |
| Isopropanol | IBI Scientific | IB15730 |
| Ethanol | Decon Laboratories | 3916EA |
| Critical commercial assays |  |  |
| NEBuilder HiFi DNA Assembly Master Max | New England Biolabs | E2621S |
| Zyppy Plasmid Miniprep Kit | Zymo Research | D4019 |
| Q5 High-Fidelity 2x Master Mix | New England Biolabs | M0492S |
| Taq 2x Master Mix | New England Biolabs | M0270L |
| NEBuilder HiFi DNA Assembly Master Max | New England Biolabs | E2621S |
| AMPure XP Beads for DNA Cleanup | Beckman Coulter | A63880 |
| PerfeCTa SYBR Green FastMix | Quantobio | 95072-250 |
| SuperScript™ III First-Strand Synthesis System | Invitrogen | 18080051 |
| DNA-free DNA Removal Kit | Invitrogen | AM1906 |
| Experimental models: Organisms/strains |  |  |
| *Magnaporthe oryzae* strain Guy11 | Mark Farman<br>(University of Kentucky) | N/A |
| *E. coli* NEB 5-alpha | New England Biolabs | C2987I |
| *Agrobacterium tumefaciens* AGL-1 | GoldBIO | CC-208-5x50 |
| Recombinant DNA |  |  |
| pMY200 plasmid | Mark Farman<br>(University of Kentucky) | N/A |
| Software and algorithms |  |  |
| FungiDB | https://fungidb.org | N/A |
| NanoDrop spectrophotometer software | Fisher Scientific | N/A |
| Other |  |  |
| Sterile Petri dishes (100mm x 15mm) | VWR | 25384-088 |
| Sterile Petri dishes (60mm x 15mm) | VWR | 25384-168 |
| VWR Bacti Cell Spreaders | VWR | 60828-680 |
| Sterile forceps | Fisher Scientific | 16-100-107 |
| 15 mL Falcon tubes | VWR | 21008-216 |
| 50 mL Falcon tubes | Fisher Scientific | 14-375-150 |
| 1.7 mL microcentrifuge tubes | MedSupply Partners | 15-1151 |
| P200 Pipette tips | VWR | 76323-390 |
| P1000 pipette tips | VWR | 613-6419 |
| Hemocytometer | Fisher Scientific | 02-671-51B |
| Miracloth | EMD Millipore Corp | 475855-1R |
| Whatman Filter Paper | The Lab Depot | 28462-025-PK |
| Growth chamber | Geneva Scientific | 1-36NL |
| Shaker incubator | Fisher Scientific | 07-202-157 |
| Biosafety cabinet | NUAIRE | NU-540 |
| Mastercycler Nexus Thermal Cyclers | Eppendorf | EP6348000029 |
| Agarose gel electrophoresis system | BIO-RAD | 1704466 |
| Electrophoresis Power Supply | Mashall Scientific | FS-FB300 |
| Agilent 2100 Bioanalyzer | Agilent Technologies, Inc. | G2939A |
| CFX96 Touch Real-Time PCR Detection System | BIO-RAD | CFX96 |
| Qubit 4 Fluorometer | Invitrogen | Q33238 |
| Bead mill homogenizer | BioSpec Products | 607 |
| Centrifuge (benchtop) | Fisher Scientific | 13-100-675 |

## MATERIALS AND EQUIPMENT

### Prepare medium and stock solution

#### (Timing: 1 week)

##### Oatmeal Agar (OM)

Dissolve 25 g/L Quaker Old Fashioned Oats in 400–500 mL deionized water. Warm in a microwave for 3–5 min, then add a stir bar. Heat and stir on a hot plate at 60°C for 1 hour. Strain the solution through a cheesecloth, and adjust the volume to 1 L before adding 15 g/L of bacteriological-grade agar. Mix well, then autoclave at 121°C for 20 min in a liquid cycle. After autoclaving, pour the sterilized solution into 100 × 15 mm petri dishes and store at 4°C for up to 2 months.

**Complete medium (CM)**
| Reagent | Final concentration | Amount |
| --- | --- | --- |
| Sucrose | 10 g/L | 10 g |
| Yeast extract | 6 g/L | 6 g |
| Casamino acid | 6 g/L | 6 g |
| Agar | 15 g/L | 15 g |
| ddH<sub>2</sub>O | n/a | up to 1 L |

Autoclave at 121°C for 20 min in liquid cycle.
Agar is not required for liquid CM. Store at room temperature up to 6 months.

**Induction medium**
| Component | Final concentration | Amount |
| --- | --- | --- |
| Sucrose | 10 g/L | 10 g |
| 1000X Trace Element Solution | 1 mL/L | 1 mL |
| KNO<sub>3</sub> | 7.14 g/L | 7.14 g |
| KCl | 0.52 g/L | 0.52 g |
| MgSO<sub>4</sub>·7H<sub>2</sub>O | 0.52 g/L | 0.52 g |
| KH<sub>2</sub>PO<sub>4</sub> | 1.52 g/L | 1.52 g |
| Thiamine (from 1g/L stock) | 1 mg/L | 1 mL |
| Biotin (from 50µg/mL stock) | 5 ng/L | 100 µL |
| Ampicillin (from 100mg/mL stock) | 100 µg/L | 1 mL |
| ddH<sub>2</sub>O | n/a | up to 1 L |

Adjust pH to 6.5 with NaOH. Autoclave at 121°C for 20 min in liquid cycle. After autoclaving, cool the medium to 50°C before adding filter-sterilized thiamine and biotin. Store at room temperature up to 6 months.

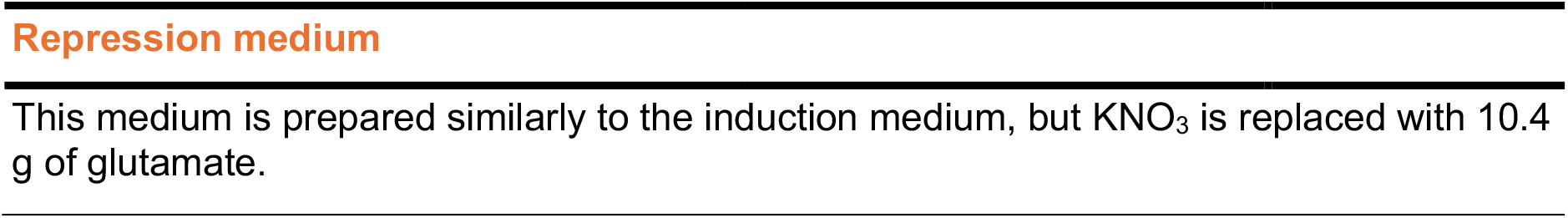

1000X Trace Elements Solution
| Reagent | Final concentration | Amount |
| --- | --- | --- |
| ZnSO<sub>4</sub> | 136.25 mM | 22 g |
| H<sub>3</sub>BO<sub>3</sub> | 177.91 mM | 11 g |
| MnCl<sub>2</sub>·4H<sub>2</sub>O | 25.26 mM | 5 g |
| FeSO<sub>4</sub>·7H<sub>2</sub>O | 17.98 mM | 5 g |
| CoCl<sub>2</sub> | 13.09 mM | 1.7 g |
| CuSO<sub>4</sub>·5H<sub>2</sub>O | 6.42 mM | 1.6 g |
| Na<sub>2</sub>MoO<sub>4</sub>·2H<sub>2</sub>O | 6.20 mM | 1.5 g |
| Disodium EDTA | 134.32 mM | 50 g |
| ddH<sub>2</sub>O | n/a | up to 1 L |

Store at room temperature up to 6 months. Precipitate may form, heat to 65°C to dissolve

Induction Medium (IM) Liquid
| Component | Final concentration | Amount |
| --- | --- | --- |
| Minimal Salts (25X stock) | 2X | 80 mL |
| 1M MES (pH 5.3) | 20 mM | 20 mL |
| 1M Glucose | 5 mM | 5 mL |
| 100% Glycerol | 0.25% (v/v) | 2.5 mL |
| Thiamine (from 1g/L stock) | 1 mg/L | 1 mL |
| Acetosyringone (from 100mM stock) | 200 µM | 2 mL |
| ddH<sub>2</sub>O | n/a | up to 1 L |

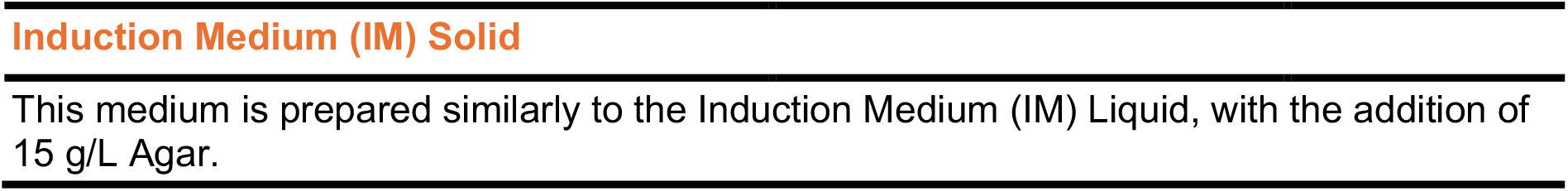

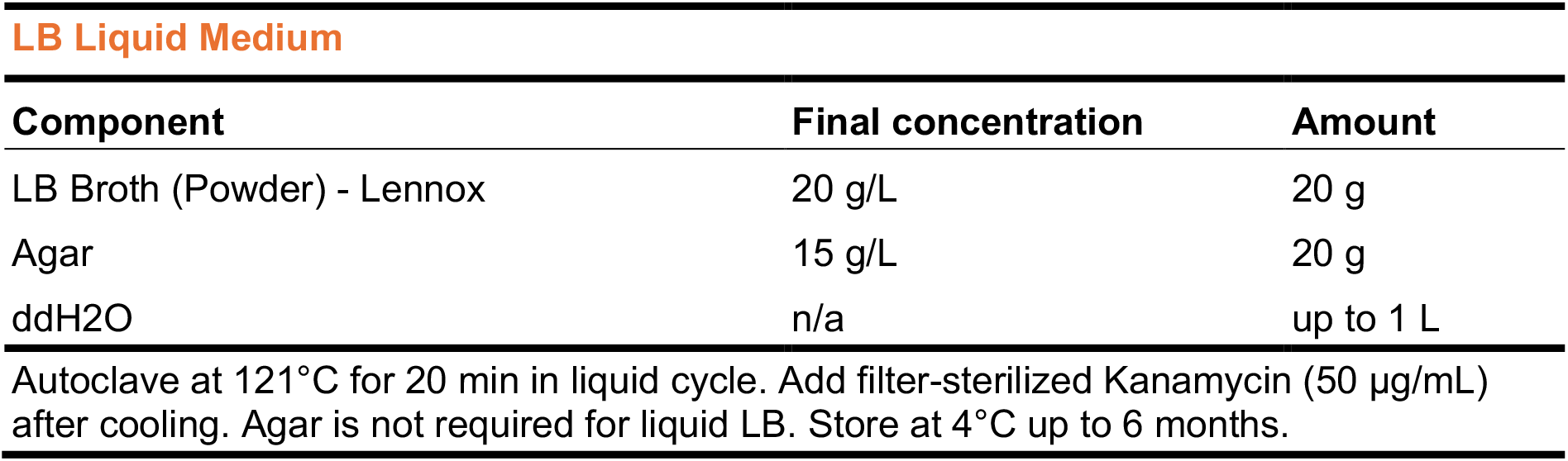

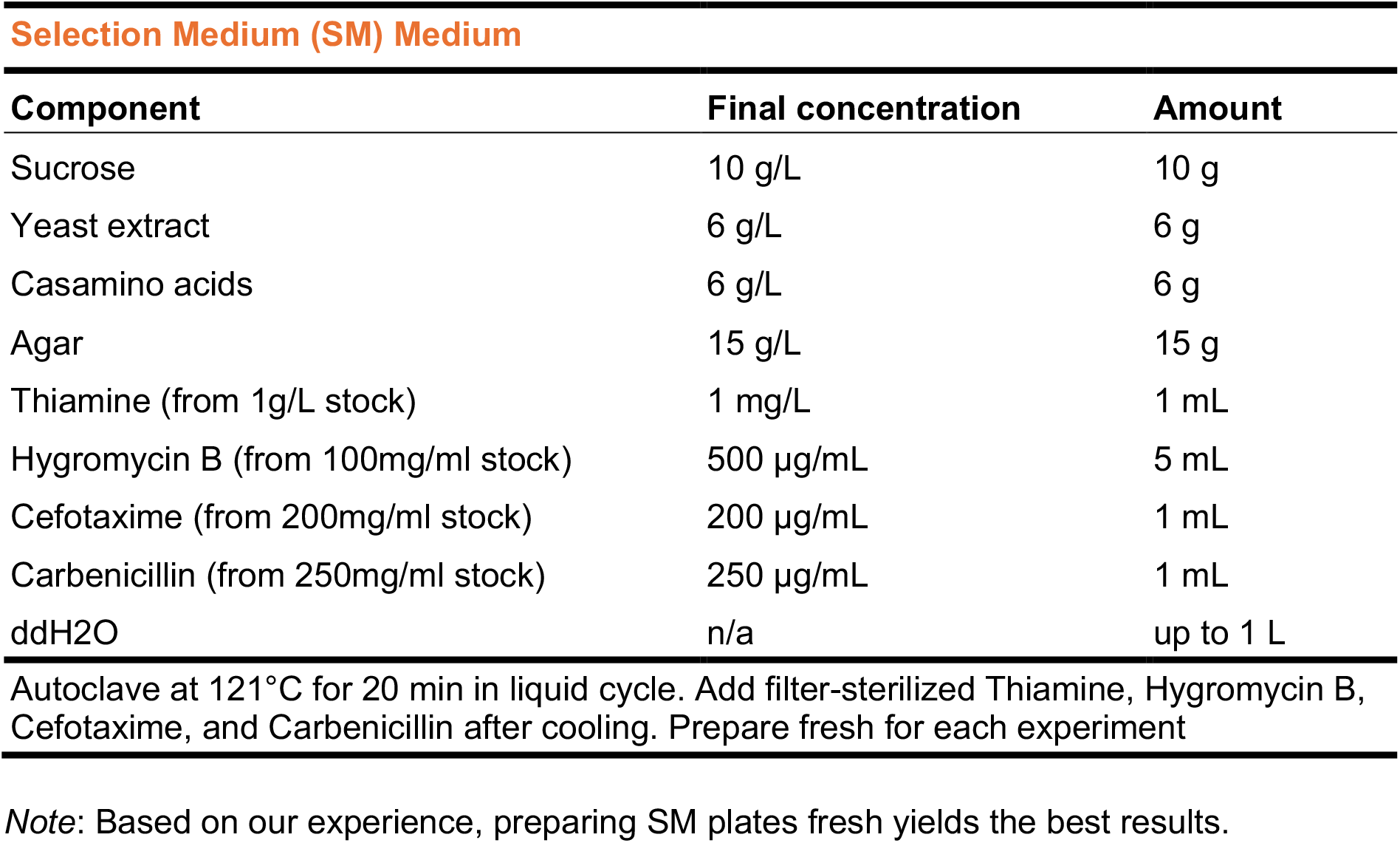

Selection Medium (SM) Medium
| Component | Final concentration | Amount |
| --- | --- | --- |
| Sucrose | 10 g/L | 10 g |
| Yeast extract | 6 g/L | 6 g |
| Casamino acids | 6 g/L | 6 g |
| Agar | 15 g/L | 15 g |
| Thiamine (from 1g/L stock) | 1 mg/L | 1 mL |
| Hygromycin B (from 100mg/ml stock) | 500 µg/mL | 5 mL |
| Cefotaxime (from 200mg/ml stock) | 200 µg/mL | 1 mL |
| Carbenicillin (from 250mg/ml stock) | 250 µg/mL | 1 mL |
| ddH<sub>2</sub>O | n/a | up to 1 L |

Autoclave at 121°C for 20 min in liquid cycle. Add filter-sterilized Thiamine, Hygromycin B, Cefotaxime, and Carbenicillin after cooling. Prepare fresh for each experiment
*Note:* Based on our experience, preparing SM plates fresh yields the best results.

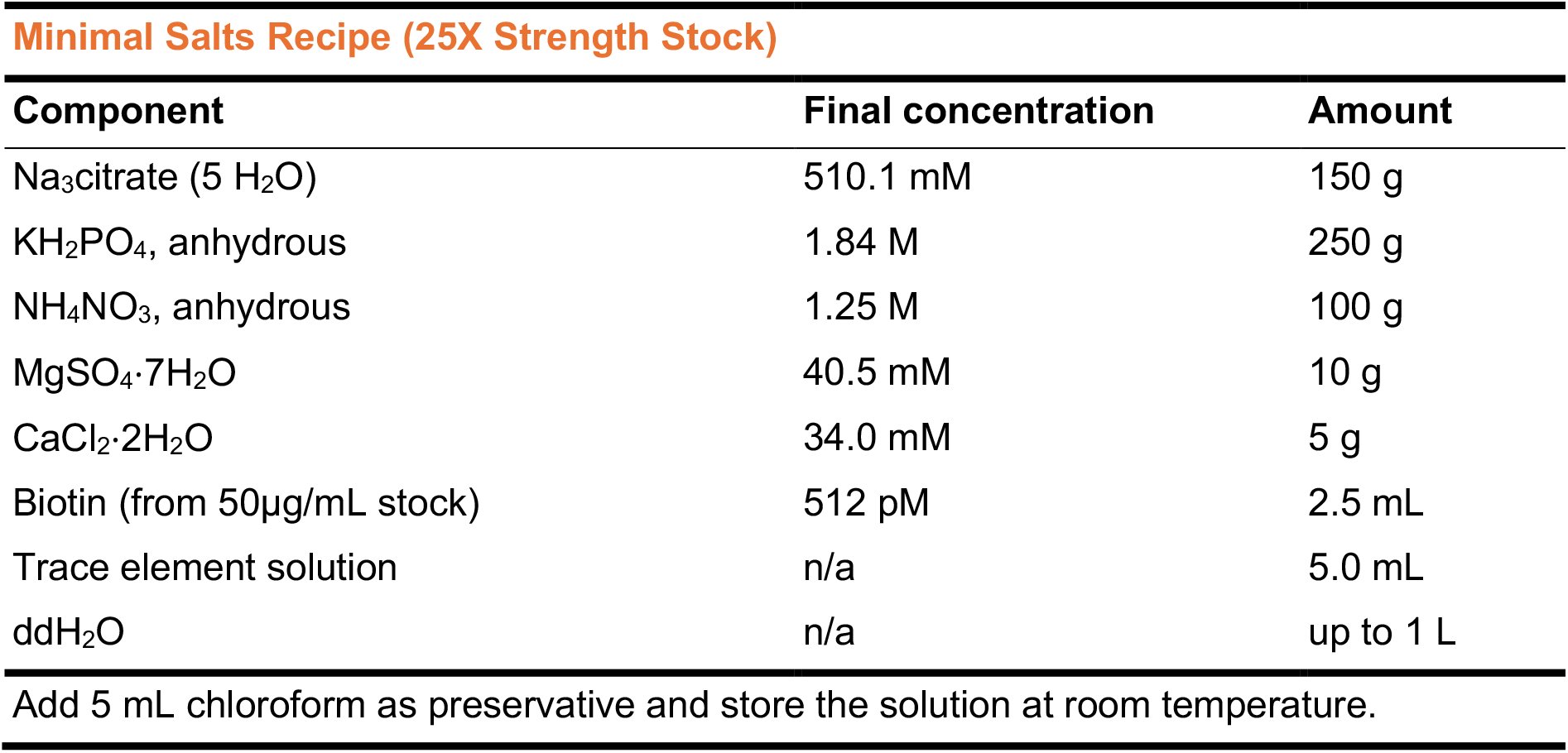

| Component | Final concentration | Amount |
| --- | --- | --- |
| Na<sub>3</sub>citrate (5 H<sub>2</sub>O) | 510.1 mM | 150 g |
| KH<sub>2</sub>PO<sub>4</sub>, anhydrous | 1.84 M | 250 g |
| NH<sub>4</sub>NO<sub>3</sub>, anhydrous | 1.25 M | 100 g |
| MgSO<sub>4</sub>·7H<sub>2</sub>O | 40.5 mM | 10 g |
| CaCl<sub>2</sub>·2H<sub>2</sub>O | 34.0 mM | 5 g |
| Biotin (from 50µg/mL stock) | 512 pM | 2.5 mL |
| Trace element solution | n/a | 5.0 mL |
| ddH<sub>2</sub>O | n/a | up to 1 L |

Add 5 mL chloroform as preservative and store the solution at room temperature.

**Trace Element Solution**
| Component | Final concentration | Amount |
| --- | --- | --- |
| Citric acid ·H<sub>2</sub>O | 237.9 mM | 5.0 g |
| ZnSO<sub>4</sub>·7H<sub>2</sub>O | 17.3 mM | 5.0 g |
| Fe(NH<sub>4</sub>)<sub>2</sub>(SO<sub>4</sub>)<sub>2</sub>·6H<sub>2</sub>O | 2.5 mM | 1.0 g |
| CuSO<sub>4</sub>·5H<sub>2</sub>O | 1.0 mM | 0.25 g |
| MnSO<sub>4</sub>·H<sub>2</sub>O | 0.3 mM | 0.05 g |
| H<sub>3</sub>BO<sub>3</sub> | 0.8 mM | 0.05 g |
| Na<sub>2</sub>MoO<sub>4</sub>·2H<sub>2</sub>O | 0.2 mM | 0.05 g |
| ddH<sub>2</sub>O | n/a | up to 100 mL |

Store at room temperature up to 6 months.

## STEP-BY-STEP METHOD DETAILS

### Design of Promoter Replacement Construct (ProRC)

#### (Timing: 2-4 weeks)

This step is required to design the DNA construct for replacing the native promoter of a target gene with a conditional promoter. To establish conditional gene expression in *Magnaporthe oryzae* strain Guy11, we targeted the *BUF1* gene (MGG_02252), which encodes a trihydroxynaphthalene reductase involved in fungal melanin biosynthesis ^4^.

Disruption of *BUF1* results in a characteristic buff (tan/orange) mycelial color, distinct from the wild-type gray/black pigmentation. We aim to place *BUF1* under the control of the *M. oryzae NIA1* (*pMoNIA1*) promoter (from MGG_06062), which is known to express conditionally based on nitrogen availability ^1^. In this step, we describe the procedure to design the specific DNA construct for this promoter replacement, which relies on homologous recombination (Figure 1).

**Figure 1.**
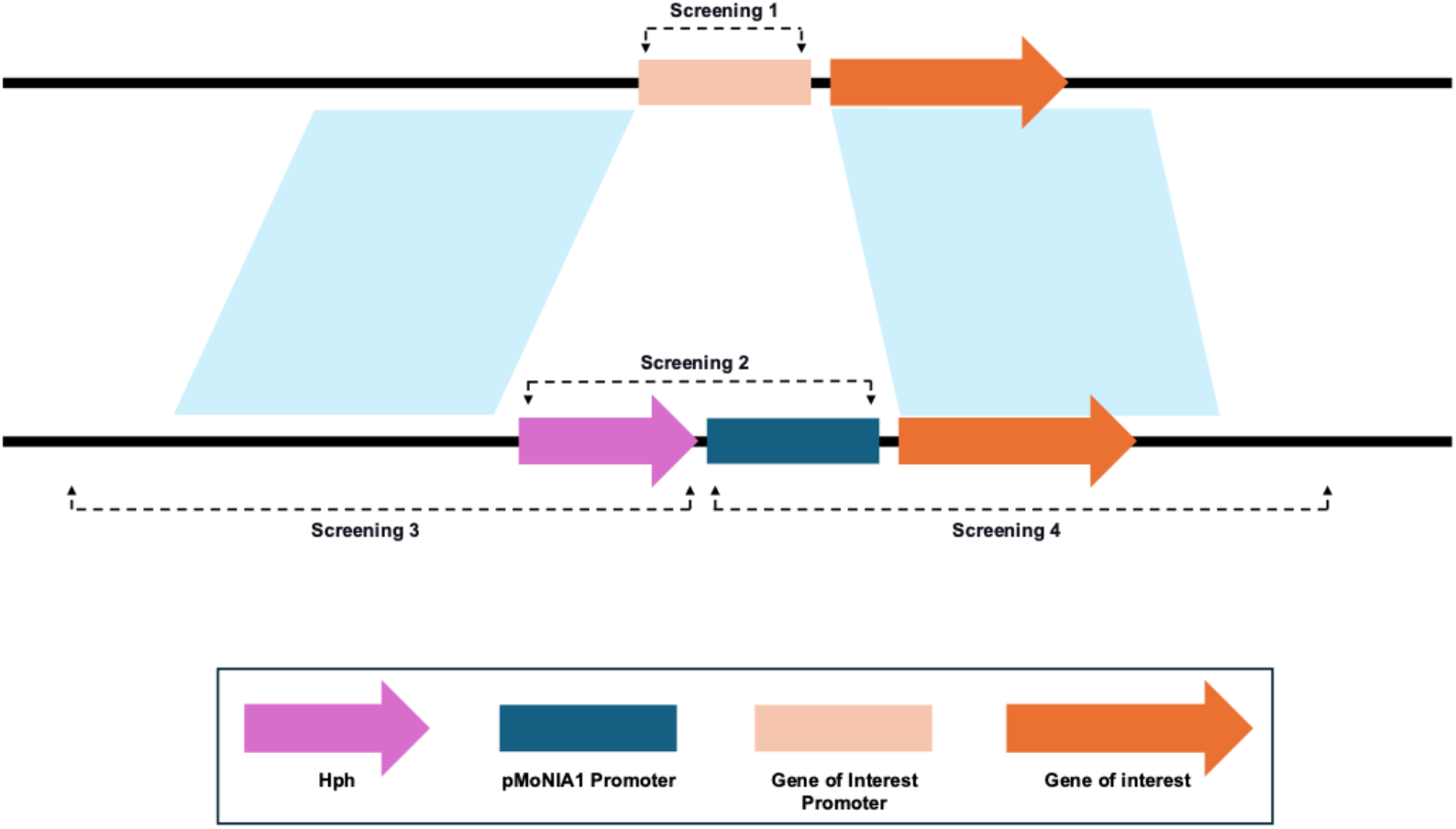
Schematic representation of the Conditional Promoter Replacement (CPR) strategy in *M. oryzae*. This schematic illustrates the targeted gene replacement method for establishing conditional gene expression. The top line represents the wild-type (WT) genomic locus, where the Gene of Interest Promoter (light orange box) precedes the Gene of Interest (orange arrow). The bottom line shows the resulting locus after a successful Homologous Recombination (HR) event. The native promoter is precisely replaced by the CPR cassette, which consists of the *pMoNIA1* Promoter (dark blue box) and the Hph (hygromycin resistance) selectable marker (pink arrow). The light blue shaded areas denote the Left and Right Flanks of homology that guide the precise replacement event. Dashed lines indicate the PCR primer binding sites used for molecular verification: Screening 1 confirms the absence of the native promoter in transformants; Screening 2 confirms the presence of the *pMoNIA1* promoter; and Screening 3 and Screening 4 are positioned outside the construct boundaries to confirm the precise integration of the entire hph-pMoNIA1 cassette at the intended locus.

1. Obtain the genomic sequence of the *BUF1* gene (MGG_02252) and its coding sequence (CDS) from a relevant fungal genome database (FungiDB: https://fungidb.org/fungidb/app).
2. Identify the native *BUF1* promoter region (507 bp) located immediately upstream of the *BUF1* CDS. This 507 bp region is the target for replacement.
3. Identify the upstream homologous region (referred to as “Left Flank” in primer design) of 1407 bp, located immediately upstream of the 507 bp native *BUF1* promoter, and the downstream homologous region (referred to as “Right Flank” in primer design) of 2728 bp, located immediately downstream of the 507 bp native *BUF1* promoter (and thus overlapping with the beginning of the *BUF1* coding sequence). These flanks will serve as homology arms for recombination (Figure 1).
4. Identify the *M. oryzae NIA1* (*pMoNIA1*) promoter sequence (1839 bp) by obtaining the upstream untranslated region (UTR) of the MGG_06062 nitrate reductase gene from FungiDB. This 1839 bp *pMoNIA1* promoter will replace the native *BUF1* promoter.
5. The hygromycin resistance gene (*hph*, 1440 bp) will be amplified from the pBShyg2 plasmid and inserted along with the *pMoNIA1* promoter. The *pMoNIA1* promoter must be positioned immediately upstream of the *BUF1* CDS to drive its conditional expression.
6. Design primers to amplify the 1407 bp Left Flank region. These primers should include overhangs homologous to the pMY200 vector backbone (after PmeI digestion) and the *hph* gene for Gibson assembly. The primers used are proBUF1_LF_F and proBUF1_LF_R2.
7. Design primers to amplify the 1440 bp *hph* gene. These primers should include overhangs homologous to the Left Flank and the *pMoNIA1* promoter for Gibson assembly. The primers used are Hyg_CE_F2 and Hyg_CE_R2.
8. Design primers to amplify the 1839 bp *pMoNIA1* promoter sequence. These primers should include overhangs homologous to the *hph* gene and the *BUF1* CDS for Gibson assembly, ensuring the *pMoNIA1* promoter is correctly positioned to drive *BUF1* expression. The primers used are pMONIA1_F2 and pMONIA1_R2.
9. Design primers to amplify the 2728 bp Right Flank region. These primers should include overhangs homologous to the *BUF1* CDS (specifically, the region immediately following the native promoter) and the pMY200 vector backbone for Gibson assembly. The primers used are proBUF1_RF_F2 and proBUF1_RF_R2. ***Note*:** Ensure all primers are designed with appropriate melting temperatures (Tm) and minimal secondary structures for efficient PCR amplification and Gibson assembly.

### Construct Assembly via Gibson Assembly and Plasmid Preparation

#### (Timing: 2-3 days)

Following the design of the promoter replacement construct, this step is required to amplify the individual DNA fragments and assemble them into the pMY200 binary plasmid using Gibson assembly. This results in a complete and verified plasmid ready for *Agrobacterium*-mediated transformation.

10 Perform PCR amplification of the following fragments: the 1407 bp Left Flank (LF), the 1440 bp *hph* gene, the 1839 bp *pMoNIA1* promoter, and the 2728 bp Right Flank (RF). Prepare the reaction master mix as follows: *Note*: Annealing temperature and extension time vary for each PCR fragment: (LF) 72°C,1:30 min, (*hph*) 72°C, 1:30 min, (*pMoNIA1*) 72°C, 1:30 min, (RF) 72°C, 2 min.
  a. Run a small aliquot of each PCR product on an agarose gel to verify the correct size and amplification efficiency.
  b. Purify all PCR products using Ampure XP beads (Beckman Coulter) to remove primers, dNTPs, and polymerases. Quantify purified DNA using a NanoDrop spectrophotometer (Thermo Scientific).
11 Linearize the pMY200 *Agrobacterium* binary plasmid with the blunt-end restriction enzyme PmeI. This digestion should occur between the Left Border (LB) T-DNA repeat and Right Border (RB) T-DNA repeat, creating a linearized backbone ready for insertion of the replacement construct. Verify complete digestion by agarose gel electrophoresis and purify the linearized vector using Ampure XP beads.
12 Perform Gibson Assembly and plasmid verification:
  a. Set up the Gibson Assembly reaction; combine equimolar amounts of the purified LF, *hph* gene, *pMoNIA1* promoter, RF, and the linearized pMY200 vector using NEBuilder HiFi DNA Assembly Master Mix (New England BioLabs). Assemble the fragments in the order pMY200 backbone - LF - *hph* - pMoNIA1 - RF - pMY200 backbone. Incubate the reaction at 50°C for 30 minutes.
  b. Transform the Gibson Assembly reaction product into chemically competent *E. coli* DH5α cells. Plate transformed cells on LB plates containing kanamycin. Incubate overnight at 37°C.
  c. Pick 20 colonies from the plate and culture them overnight in LB liquid medium supplemented with kanamycin. Isolate plasmid DNA using a Zyppy Plasmid Miniprep Kit (Zymo Research) from these cultures. Verify the successful assembly of the final plasmid, designated pMY200-ProRC, by restriction enzyme digestion and by full plasmid sequencing by Plasmidsaurus using Oxford Nanopore Technology with custom analysis and annotation.

| Reagent | Amount |
| --- | --- |
| Q5 High-Fidelity 2x Master Mix | 25 µL |
| DNA template (~ 10 ng/ µL) | 5 µL |
| forward primer (10 µM) | 2.5 µL |
| reverse primer (10 µM) | 2.5 µL |
| Molecular water | 15 µL |
| Total | 50 µL |

| PCR cycling conditions |  |  |  |
| --- | --- | --- | --- |
| Steps | Temperature | Time | Cycles |
| Initial Denaturation | 98°C | 30 s | 1 |
| Denaturation | 98°C | 10 s | 32 |
| Annealing | Variable | 30 s |  |
| Extension | 72°C | Variable |  |
| Final extension | 72°C | 2 min | 1 |
| Hold | 4°C |  |  |

*Note:* Annealing temperature and extension time vary for each PCR fragment: (LF) 72°C, 1:30 min, (*hph*) 72°C, 1:30 min, (*pMoNIA1*) 72°C, 1:30 min, (RF) 72°C, 2 min.

### *Agrobacterium tumefaciens*-Mediated Transformation (ATMT) of *Magnaporthe oryzae*

#### (Timing: 3-4 weeks)

This step describes the transformation of *M. oryzae* using the *Agrobacterium tumefaciens*-mediated transformation (ATMT) method ^5^. The plasmid pMY200-ProRC, which contains the LF-*hph*-pMoNIA1-RF replacement cassette, is introduced into the fungal genome via co-cultivation with *A. tumefaciens*.

13 Prepare *A. tumefaciens*:
  a. Gently mix 1 µL of 20 ng/µL pMY200-ProRC plasmid DNA with 50 µL of electrocompetent *A. tumefaciens* strain AGL-1 cells in a 1.7 mL microcentrifuge tube on ice by flicking the tube.
  b. Transfer the cell/DNA mixture into a pre-chilled 0.1 cm electroporation cuvette. Apply a single pulse of electroporation (2.5 kV, 25 µF, 200 Ω) with a typical time constant of 4-5 ms.
  c. Immediately add 1 mL of warm liquid LB medium to the cuvette and transfer the entire suspension to a 15 mL Falcon tube. Incubate with shaking at 28 °C for 4 hours to allow recovery.
  d. Plate 100 µL aliquots of the recovered culture on solid LB medium supplemented with 100 µg/mL kanamycin and incubate at room temperature for 2 days.
  e. Pick a single colony from the transformation plate and inoculate it into liquid LB medium supplemented with 100 µg/mL kanamycin. Grow overnight at 28°C with shaking. Prepare a glycerol stock from this overnight culture (mix 600 µL of culture with 600 µL of 50% glycerol) and store at −80°C.
  f. For subsequent co-cultivation experiments, take a loopful smear from the frozen glycerol stock and streak it onto a fresh solid LB medium plate supplemented with 100 µg/mL kanamycin. Incubate at 28°C until actively growing colonies are visible (typically 2 days). This plate will be used directly for the follow up co-cultivation step.
14 Prepare *M. oryzae* spore suspension:
  a. Activate *M. oryzae* strain Guy11 by transferring a stored filter paper from −20°C to a CM plate and incubating at 25°C under continuous light for 6 days. Then, transfer a 1 cm square from this plate to an Oatmeal Agar (OM) plate and grow under continuous light at 25°C for 30 days to allow sporulation.
  b. Harvest spores by flooding the plate with 5-10 mL of sterile water and gently scraping the surface with a sterile bacterial spreader. Filter the resulting spore suspension through 0.2 µm Miracloth into a 15 mL sterile conical tube.
  c. Concentrate the spores by centrifuging at 5700 rpm for 5 minutes and remove excess water. Count spores using a hemocytometer and dilute to a final concentration of 5.0×10^5^ spores/mL using sterile water.
15 Perform co-cultivation:
  a. Prepare two co-cultivation agar plates (IM solid medium, containing 5mM glucose and 200 µM AS) and cover each with autoclaved black filter paper squares (1cm X 1cm).
  b. Label the plates with date, *Agrobacterium* strain, fungal strain, and plate number.
  c. Vigorously mix the spore suspension and dispense 100 µL onto each plate, distributing small droplets evenly.
  d. Using a P1000 pipette tip, scoop a visible mass of *A. tumefaciens* and dot it evenly across the plate surface.
  e. Using a sterile bacterial spreader (flat side), gently but firmly massage the plate to mix the spores and bacteria thoroughly. Continue until the *Agrobacterium* is no longer visibly distinct.
  f. Place the plates right side up on a lighted shelf at 25°C under continuous light.
16 Perform bacterial elimination and selection of transformants:
  a. After 48 hours, use sterile forceps to transfer 8–10 filter paper squares from each co-cultivation plate onto selection plates (CM solid medium). Place filter papers with at least 0.5 cm of space between them to allow resistant transformants to grow out. **Critical:** Do not move the paper after placement to avoid accidental overgrowth of the fungi.
  b. Ensure the selection medium contains 300 µg/mL hygromycin, 200 µg/mL cefotaxime, and 250 µg/mL carbenicillin.
  c. Incubate selection plates at 25°C under continuous light for 3-8 days until resistant fungal colonies begin growing off of the placed filter paper.
17 Collect transformants:
  a. Pick individual hygromycin-resistant mycelia that have grown out from the filter paper onto the selection plate using a sterile scalpel.
  b. Transfer each picked mycelia to fresh CM Agar plates supplemented with 300 µg/mL hygromycin to ensure stability and purity.

### Screening of Transformants

#### (Timing: 3-5 days)

This step confirms the successful replacement of the native *BUF1* promoter with the *hph*-*pMoNIA1* cassette.

18 Perform fungal DNA extraction:
  a. Grow selected transformants in liquid CM culture for 3-5 days.
  b. Harvest mycelia by filtration and extract genomic DNA from the mycelia using a rapid DNA extraction method ^6^. Quantify DNA using a NanoDrop spectrophotometer.
19 Perform PCR verification of promoter replacement:
  a. Design PCR primers to specifically amplify:
    i. A region within the native *BUF1* promoter (to confirm its absence in successful transformants). The primer pair is Screening1-F and Screening1-R, which should not yield a band in successful transformants but is 435 bp in the Wild Type (WT).
    ii. A region spanning the whole *hph*-pMoNIA1 cassette. The primer pair is Screening2-F and Screening2-R, yielding an expected size of 3,243 bp.
    iii. A region spanning the left side of the *hph*-pMoNIA1 cassette (from outside the construct). The primer pair is Screening3-F and Screening3-R, yielding an expected size of 3,342 bp. **Note:** This primer pair goes outside of the construct to ensure it is sitting in the correct position.
    iv. A region spanning the right side of the KO placement (from outside the construct). The primer pair is Screening4-F and Screening4-R, yielding an expected size of 6,516 bp. **Note:** This primer pair goes outside of the construct to ensure it is sitting in the correct position.
  b. Perform PCR reactions using the extracted genomic DNA from transformants as a template, as indicated below
  c. Analyze PCR products by agarose gel electrophoresis to confirm the expected band sizes for successful promoter replacement and the absence of the native *BUF1* promoter.

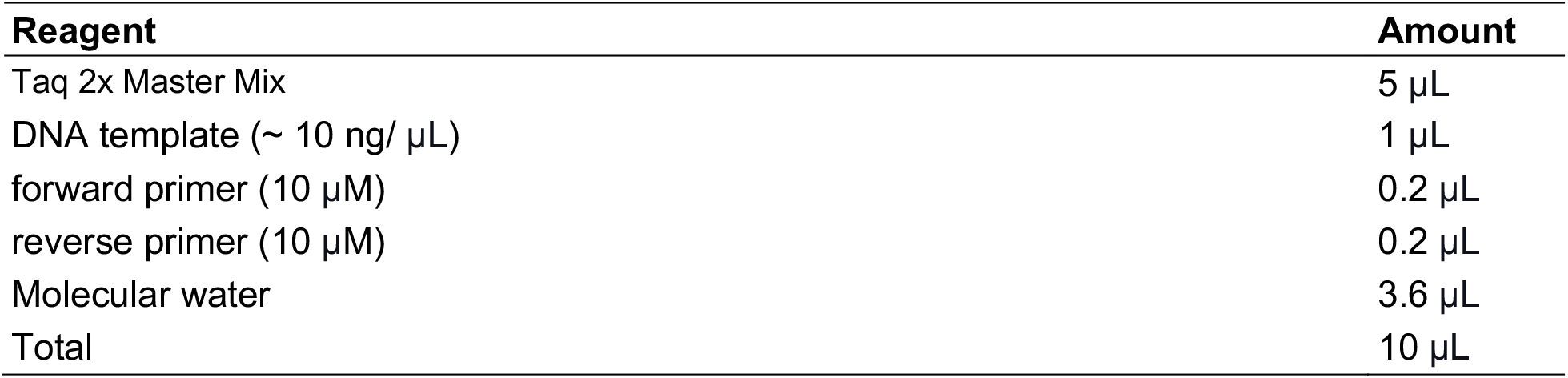

| Reagent | Amount |
| --- | --- |
| Taq 2x Master Mix | 5 µL |
| DNA template (~ 10 ng/ µL) | 1 µL |
| forward primer (10 µM) | 0.2 µL |
| reverse primer (10 µM) | 0.2 µL |
| Molecular water | 3.6 µL |
| Total | 10 µL |

| **PCR cycling conditions** |  |  |  |
| --- | --- | --- | --- |
| Steps | Temperature | Time | Cycles |
| Initial Denaturation | 95°C | 30 s | 1 |
| Denaturation | 95°C | 30 s | 30 |
| Annealing | Variable | 60 s |  |
| Extension | 68°C | Variable |  |
| Final extension | 68°C | 5 min | 1 |
| Hold | 4°C |  |  |

*Note:* Annealing temperature and extension time vary for each PCR reaction: (i) 59°C, 30 sec, (ii) 63°C, 3:30 min, (iii) 60°C, 3:30 min, (iv) 60°C, 6:30 min.

### Phenotypic Analysis of Conditional BUF1 Expression

#### (Timing: 5–7 days)

This step provides the initial visual validation that the promoter is correctly controlling expression in a nitrogen-dependent manner, correlating the molecular integration confirmed in Step 19 with the expected color switch.

20 Grow transformants (T1 and T2) confirmed to have correct promoter replacement (Step 19) by transferring a small mycelial plug onto two different solid media plates:
  a. Inducing Medium: Minimal Medium containing 7.14 g/L KNO_3_ (Inducing Condition).
  b. Repressing Medium: Minimal Medium containing 10.4 g/L glutamate (Repressing Condition).
21 Include the wild-type *M. oryzae* as a control on both media types.
22 Incubate all plates at 25°C under continuous light for 5–7 days.
23 Observe and document the resulting colony pigmentation (Figure 3).

### Quantification of Conditional *BUF1* Expression via qRT-PCR

#### (Timing: 2-5 days)

This step validates the conditional gene expression system by quantifying the relative transcription levels of the targeted gene (*BUF1*) in transformants grown under inducing (NO_3_^−^) and repressing (Glutamate) conditions. This analysis requires the use of two stably expressed reference genes for accurate normalization.

24 Determine Fungal Growth Conditions and Harvest Mycelium
  a. Grow three independent biological replicate cultures of purified *M. oryzae* pMoNIA1::*BUF1* transformants and wild-type strain Guy11 as a control in agitated liquid Induction Medium (containing 7.14 g/L KNO_3_) and Repression Medium (containing 10.4 g/L glutamate) for 5 days at 25°C with agitation (180 RPM).
  b. Harvest fungal mycelium from each replicate by filtration (through Miracloth). Wash mycelia briefly with sterile water to remove residual media. Dry the fungal mycelium with paper towels, then immediately snap-freeze approximately 100 mg of the fungal biomass in liquid nitrogen and store it at −80°C prior to RNA extraction. CRITICAL: Rapid freezing in liquid nitrogen is essential to prevent RNA degradation. Process samples one at a time to minimize time between harvest and freezing.
25 Isolate Total RNA and Synthesize cDNA
  a. Extract total RNA from the frozen fungal mycelium using the TRIzol® Reagent method (Invitrogen) following the manufacturer’s protocol. Note: Grind frozen mycelium to a fine powder in a pre-chilled mortar and pestle with liquid nitrogen before adding TRIzol to maximize RNA yield and quality.
  b. Eliminate potential genomic DNA contamination from the total RNA sample using a DNA-free kit according to the manufacturer’s instructions.
  c. Quantify the resulting RNA concentration using a Qubit 4 fluorometer (Invitrogen). Normalize RNA samples to a uniform concentration (e.g., 500-1000 ng/μL). Note: Fluorometric quantification (e.g., Qubit) is generally preferred over spectrophotometry (e.g., NanoDrop) for determining RNA concentration prior to cDNA synthesis, as it is highly specific to nucleic acids and avoids overestimation caused by non-nucleic acid contaminants (e.g., salts, phenol, or proteins) often present in crude extracts.
  d. Verify the integrity of the total RNA for each of the three replicates using the Agilent 2100 Bioanalyzer Instrument. CRITICAL: Ensure that the RNA Integrity Number (RIN) for pure fungal RNA samples (vegetatively grown tissues) is ≥7.5 to ensure minimal degradation.
  e. Use a standardized amount of isolated total RNA (e.g., 1.0 μg) from each of the three independent biological replicates to synthesize complementary DNA (cDNA) using SuperScript™ III First-Strand Synthesis System (Invitrogen). Note: Prepare a minus-Reverse Transcriptase (−RT) control for each sample to check for genomic DNA contamination. Use the same amount of RNA as experimental samples but omit reverse transcriptase from the reaction.
26 Perform Quantitative Real-Time PCR (qRT-PCR)
  a. Design and synthesize primer pairs specific for the target gene (*BUF1*) and the two validated internal reference genes for *M. oryzae* vegetative tissue: MGG_Ef1 (MGG_03641) and MGG_40s (MGG_02872) ^7^. Primer sequences are listed in Table S1. CRITICAL: Design qRT-PCR primers to span exon-exon junctions when possible to avoid amplification of residual genomic DNA. Amplicon sizes should be 80-200 bp for optimal qRT-PCR efficiency.
  b. Test all primer pairs using standard curves (serial dilutions of pooled cDNA: 1:1, 1:5, 1:25, 1:125, 1:625) to confirm amplification efficiencies ranging from 95% to 105% (E = 1.95-2.05, corresponding to slope values between −3.1 and −3.6) and verify the absence of non-specific amplicons via melt curve analysis ^7^.
  c. Prepare the Real-Time PCR reactions using PerfeCTa SYBR Green FastMix (Quantabio) for SYBR-Assay. Note: Utilize three technical replicates per biological replicate reaction to account for pipetting variability.
  d. Introduce diluted cDNA template (e.g., 2.0 μl of 10x dilution) into the final reaction volume (20 μl). Note: Optimal cDNA dilution should be determined empirically to achieve Ct values in the range of 20-30 cycles. Include no-template controls (NTC) and −RT controls on each plate.
  e. Run the reactions using CFX96 Real-Time System (Bio-Rad) with the following thermal profile: 50°C for 2 min, followed by 95°C for 2 min; then 95°C for 15 seconds, followed by 60°C annealing for 1 min, repeating for 40 cycles.
  f. Perform a dissociation (melt) curve analysis at the end of the reaction (65°C to 95°C, 0.5°C increments) to confirm the presence of a single, specific PCR product. CRITICAL: A single sharp peak in the melt curve indicates specific amplification. Multiple peaks or broad peaks suggest primer-dimer formation or non-specific amplification; in such cases, redesign primers or optimize reaction conditions.
27 Analyze Relative Gene Expression
  a. Calculate the relative expression levels of *BUF1* using the 2 ^−ΔΔC(t)^ method ^8^. Note: ΔCt = Ct (*BUF1*) – Ct (geometric mean of reference genes); ΔΔCt = ΔCt (treatment) – ΔCt (calibrator). Use wild type grown on inducing medium as the calibrator sample.
  b. Normalize the *BUF1* expression values against the geometric mean of the expression levels of the two stable internal reference genes, MGG_Ef1α and MGG_40S ^9^. CRITICAL: The minimum number of reference genes required for accurate and reliable normalization in *M. oryzae* is two. The use of both MGG_Ef1α and MGG_40S provides greater stability for vegetative tissues than a single reference gene.
  c. Determine the fold-change in *BUF1* expression in pMoNIA1::*BUF1* transformants under repressing conditions (glutamate) compared to inducing conditions (nitrate) and to wild-type controls under both conditions.
  d. Calculate means and standard errors (SEM) from gene expression values obtained from the three independent biological replicates. Perform statistical analysis using Student’s t-test to compare expression levels across conditions. Evaluate the significance of up/down regulation with P < 0.05 considered statistically significant. Present data as bar graphs with error bars representing SEM.

## EXPECTED OUTCOMES

Successful transformation and promoter replacement are expected to yield *M. oryzae* strain Guy11 transformants where *BUF1* gene expression is tightly regulated by the nitrogen source. Transformation efficiency typically ranges from 10 to 20 transformants per co-cultivation plate. Upon initial selection on hygromycin-containing medium, resistant colonies will emerge from filter paper squares within 3–8 days of incubation at 25°C under continuous light. Approximately 10–15% of hygromycin-resistant transformants isolated from selection plates are expected to show correct integration of the hph-pMoNIA1 cassette at the native *BUF1* locus when screened by PCR genotyping. (Note: This percentage reflects the HR efficiency observed in *M. oryzae*, where random ectopic integration is highly prevalent.) The remaining transformants may represent ectopic integrations or incomplete recombination events and should be excluded from further analysis.

### Molecular validation

Genotyping by PCR should confirm precise promoter replacement and the absence of the native *BUF1* promoter (Figure 2). Using the four primer pairs described in the screening section, successful transformants will display the following characteristic PCR amplification patterns: (1) absence of the 435 bp band specific to the native *BUF1* promoter (Screening1), which should amplify only in wild-type controls, confirming complete replacement of the native promoter in transformants; (2) presence of a 3,243 bp band (Screening2) spanning the entire hph-pMoNIA1 cassette, confirming successful integration of both the hygromycin resistance marker and the conditional promoter; (3) amplification of a 3,342 bp band (Screening3) using primers positioned outside the left boundary of the construct, verifying correct integration at the 5’ junction; and (4) amplification of a 6,516 bp band (Screening4) using primers positioned outside the right boundary of the construct, confirming proper integration at the 3’ junction (Figure 3). The Screening3 and Screening4 primer pairs are particularly critical as they extend beyond the construct boundaries, providing definitive evidence that the hph-pMoNIA1 cassette integrated precisely at the intended *BUF1* locus rather than at ectopic sites. Wild-type controls should only amplify the native promoter fragment (435 bp from Screening1) and will fail to produce bands with Screening2, 3, and 4 primer pairs, while transformants with correct integration will show all three construct-specific bands (3,243 bp, 3,342 bp, and 6,516 bp) and lack the native promoter band.

**Figure 2.**
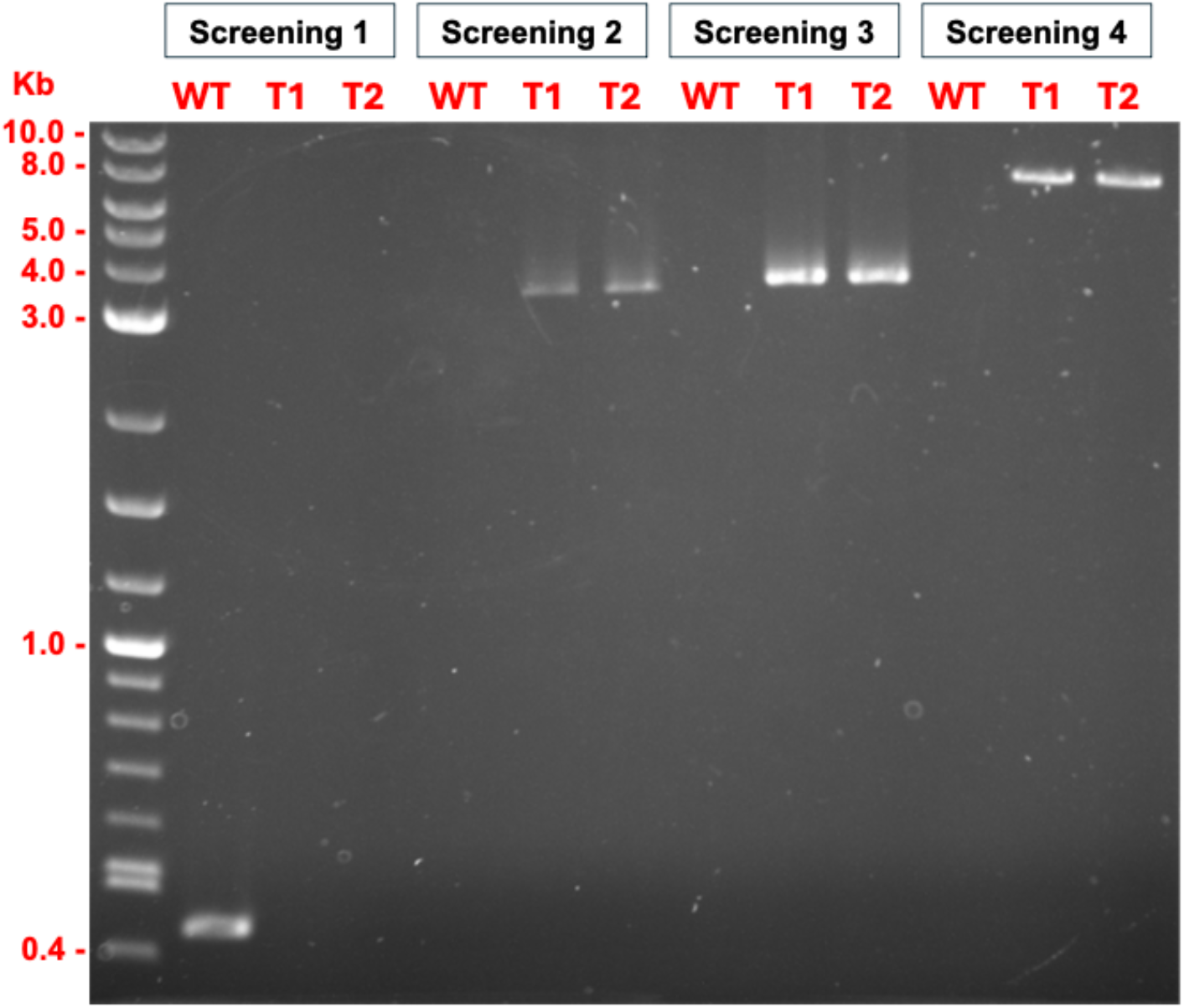
Molecular validation of pMoNIA1::*BUF1* promoter replacement by PCR genotyping. Agarose gel showing diagnostic PCR assays on wild-type (WT) *M. oryzae* and two independent transformant lines (T1 and T2). The four PCR assays demonstrate the success of the homologous recombination event. Screening 1 (Native Promoter): Shows a band only in WT (435 bp), confirming the promoter’s loss in transformants. Screening 2 (Cassette Presence): Verifies the entire hph-pMoNIA1 cassette is present in T1 and T2 (3,243 bp). Screening 3 (5’ Junction): Confirms integration at the 5’ locus border (3,342 bp). Screening 4 (3’ Junction): Confirms integration at the 3’ locus border (6,516 bp). The results confirm the absence of the native promoter band in transformants and the precise integration of the cassette at both the 5’ and 3’ junctions.

**Figure 3.**
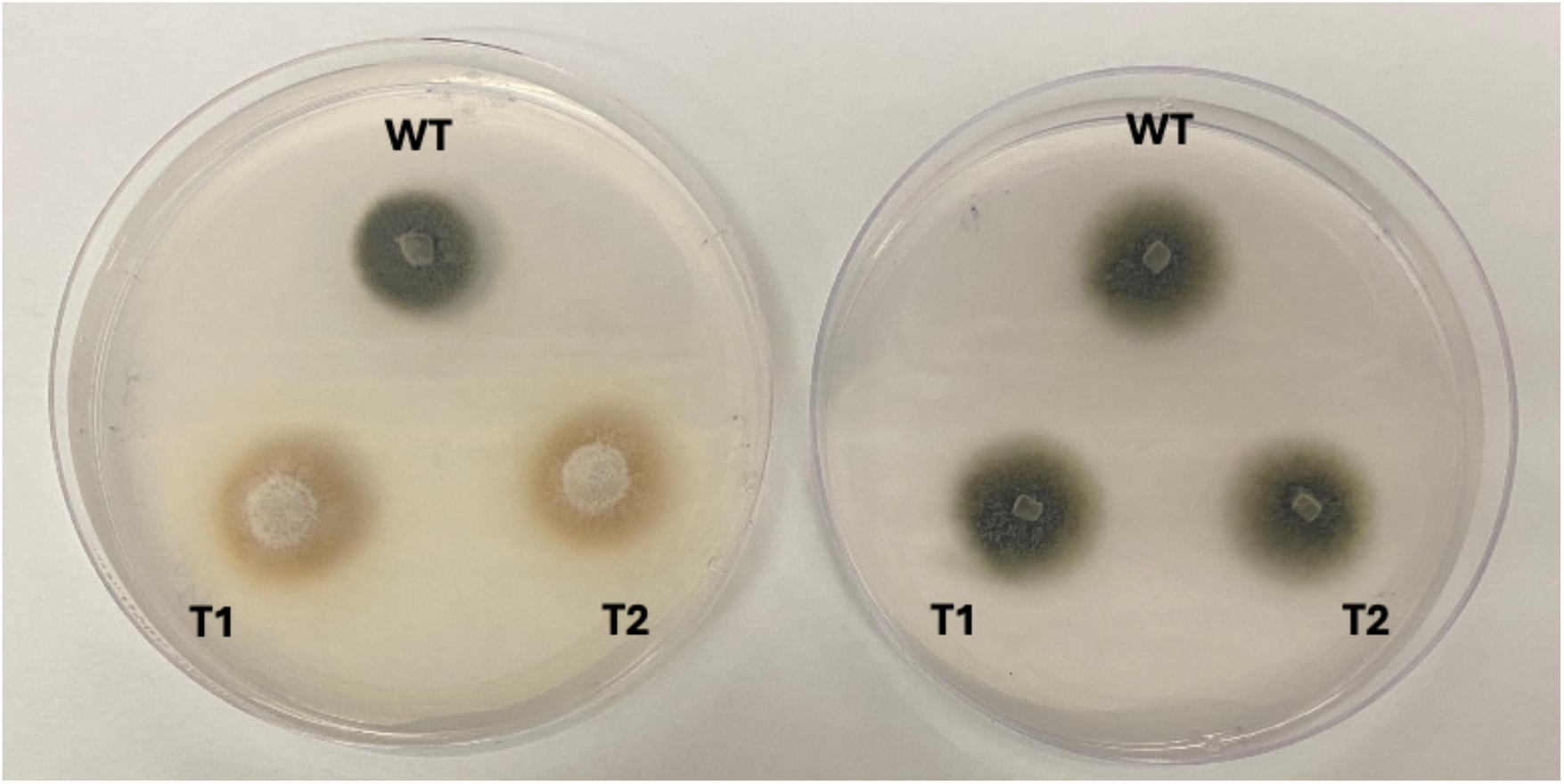
Nitrogen-dependent phenotypic switching in pMoNIA1*::BUF1* transformants. Phenotypic differences of wild-type (WT) and pMoNIA1*::BUF1* transformants (T1 and T2) grown under nitrogen-inducing and repressing conditions. Colonies were cultured on inducing medium (KNO_3_; right) or repressing medium (glutamate; left) for seven days. Transformants exhibit nitrogen source– dependent pigmentation and growth patterns, indicating transcriptional control of *BUF1* under the nitrate-inducible promoter *pMoNIA1*.

### Phenotypic validation

Transformants with verified promoter replacement should exhibit clear, nitrogen source-dependent phenotypic changes (Figure 3). Under inducing conditions (minimal medium ^10^ containing 7.14 g/L KNO_3_ as the sole nitrogen source), pMoNIA1::*BUF1* transformants should display wild-type dark gray to black pigmentation, indistinguishable from the parental strain Guy11, indicating active *BUF1* expression and functional melanin biosynthesis (Figure 2, left panels). This pigmentation should be evident within 5–7 days of growth on the inducing medium at 25°C. Conversely, under repressing conditions (minimal medium containing 10.4 g/L glutamate), transformants should display the characteristic buff, tan, or orange coloration typical of buf1 deletion mutants, confirming effective transcriptional repression of *BUF1* (Figure 2, right panels). The color transition is typically observable within 5–7 days after transfer to repressing medium. The phenotype should be fully reversible: transferring colonies from repressing back to inducing medium should restore dark pigmentation within 3–5 days, demonstrating the conditional and reversible nature of the system. As shown in Figure 2, wild-type strain Guy11 maintains dark pigmentation under both inducing and repressing conditions, while pMoNIA1::*BUF1* transformants exhibit nitrogen-dependent color switching.

### Transcriptional validation by qRT-PCR

Quantitative RT-PCR analysis should confirm that the observed phenotypic changes correlate with transcriptional regulation of *BUF1* (Figure 4, Table S2). In pMoNIA1::*BUF1* transformants grown under inducing conditions (7.14 g/L KNO_3_), *BUF1* mRNA levels are expected to range from 87.9% (T1) to 91.7% (T2) of wild-type levels (P > 0.05), confirming that the *pMoNIA1* promoter effectively restores functional *BUF1* expression to near wild-type levels. Under repressing conditions (10.4 g/L glutamate), *BUF1* transcript levels in transformants should be reduced to approximately 1.4-2.0% of wild-type induced levels, demonstrating approximately 50-fold repression (44-65-fold range, P < 0.001). This level of repression, which substantially exceeds the 18-fold down-regulation observed for the *pMoNIA1* promoter driving an EGFP reporter in *Z. tritici* ^1^, confirms the tight control achieved in the native *M. oryzae* host. Wild-type strain should maintain relatively stable *BUF1* expression across both nitrogen sources, as the native *BUF1* promoter is not nitrogen-responsive. The degree of repression may vary slightly among independent transformant lines due to positional effects, but all lines with correct integration should show substantial downregulation under repressing conditions. This molecular validation confirms that *pMoNIA1* functions as a strong, nitrogen-responsive conditional promoter in *M. oryzae* and that phenotypic changes directly reflect transcriptional regulation rather than post-transcriptional or metabolic effects.

**Figure 4.**
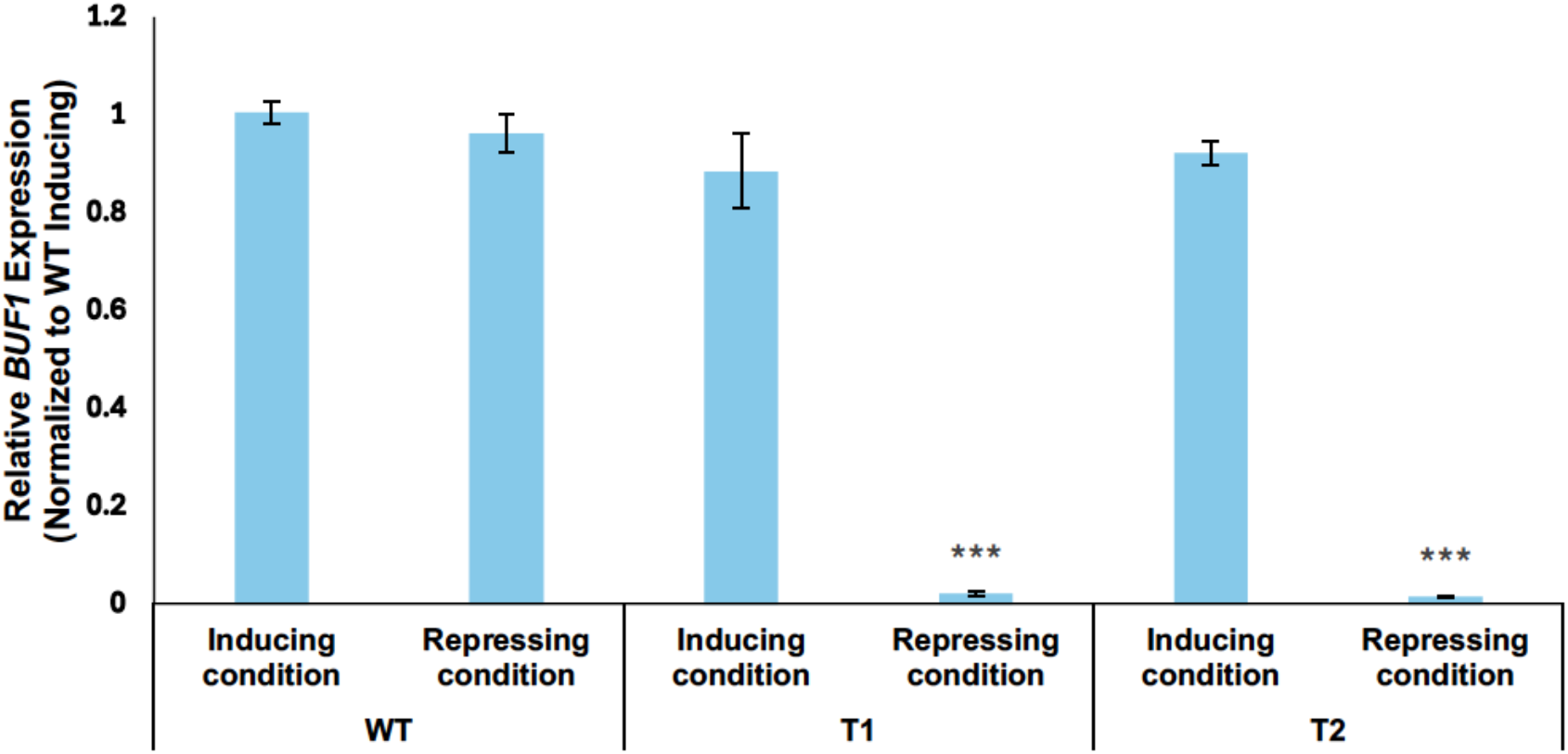
Transcriptional validation of conditional BUF1 expression by qRT-PCR. Relative *BUF1* mRNA levels in wild-type *M. oryzae* and conditional transformants (T1 and T2) under nitrogen-dependent regulation. *BUF1* mRNA levels were quantified in wild-type *M. oryzae* strain Guy11 and two independent pMoNIA1::*BUF1* transformants (T1 and T2). Expression was measured across Inducing conditions and Repressing conditions. Expression values were normalized to the geometric mean of MGGEf1α and MGG40s and presented relative to the wild-type strain under inducing conditions (set to 1.0). Data represent means ±SEM (n=3 independent biological replicates with three technical replicates each). Statistical significance was determined by paired Student’s t-test comparing the inducing vs. repressing conditions within each transformant. Significance is indicated by asterisks above the bars: *** P < 0.001. No marker is placed on the graph when the difference is not significant (P > 0.05).

## LIMITATIONS

This protocol was specifically developed and optimized for *M. oryzae* strain Guy11 and may require optimization for other *M. oryzae* isolates or related fungal species. Transformation efficiency is highly dependent on the quality of starting materials (Agrobacterium cultures and *M. oryzae* spores) and precise execution of co-cultivation conditions, which can introduce variability between experiments. While the *pMoNIA1* promoter provides tight regulation for *BUF1*, with clear phenotypic switching between inducing and repressing conditions, the degree of repression may vary when applied to other genes of interest, particularly those with different native expression levels or regulatory requirements.

The protocol requires careful preparation and quality control of all media, as trace contamination of the nitrogen source can compromise *pMoNIA1* regulation. Off-target integrations can occur despite homologous recombination, necessitating rigorous molecular screening of multiple independent transformants. The *BUF1*-based visual screening described here provides proof-of-concept validation, but application to other genes will require appropriate molecular or biochemical assays to confirm conditional expression, as most genes will not produce easily observable color changes. Researchers should also consider potential compensatory mechanisms or pleiotropic effects when essential genes are conditionally repressed, which may complicate phenotypic interpretation.

## TROUBLESHOOTING

### Problem 1

No or Very Few Transformants After Selection (Step 16)

### Potential solution

- Verify the viability of both Agrobacterium and *M. oryzae* spores before starting co-cultivation. Old or stressed cultures significantly reduce transformation efficiency.
- Confirm that the pMY200-ProRC plasmid was successfully integrated into Agrobacterium strain AGL-1 by PCR or restriction digest before use.
- Check the hygromycin concentration in selection medium. Too high concentration (>500 µg/mL) may prevent even resistant transformants from growing. Prepare fresh selection medium with freshly added antibiotics.
- Ensure proper co-cultivation conditions: verify that induction medium contains 200 µM acetosyringone and that co-cultivation occurs at 25°C for exactly 48 hours.
- Verify that bacterial growth is not overwhelming fungal growth during co-cultivation. If excessive bacterial growth is observed, reduce the amount of Agrobacterium used for co-cultivation.
- Increase the number of co-cultivation plates (3–4 plates instead of 2) to improve overall transformant yield.

### Problem 2

Bacterial Contamination Persists After Antibiotic Treatment (Step 16)

### Potential solution

- Increase cefotaxime concentration to 250–300 µg/mL and carbenicillin to 300 µg/mL in selection medium if bacterial contamination persists.
- Transfer filter paper squares with emerging fungal colonies to fresh selection plates with antibiotics every 3–4 days to continuously suppress bacterial growth.
- Once hygromycin-resistant colonies are visible growing off the filter paper, immediately transfer them to fresh CM plates with hygromycin and antibiotics to purify away from residual bacteria.
- Ensure antibiotics are freshly prepared and filter sterilized. Old or improperly stored antibiotic solutions lose effectiveness.
- If contamination persists, subculture transformants by transferring small hyphal tips (not entire colonies) to fresh medium.

### Problem 3

No Pigmentation in Transformants Under Inducing Conditions (related to Expected Outcomes)

### Potential solution

- Verify that inducing medium contains 7.14 g/L KNO_3_ as the sole nitrogen source. Insufficient nitrate will fail to induce *pMoNIA1*.
- Confirm by PCR that the *BUF1* coding sequence remains intact and in-frame with the *pMoNIA1* promoter. Frameshift mutations or deletions during construct assembly would prevent functional *BUF1* expression.
- Check the viability and growth rate of transformants. Slow-growing or unhealthy cultures may not produce visible pigmentation within the expected timeframe.
- Extend the incubation period to 10–14 days on inducing medium, as some transformants may exhibit delayed pigmentation.
- Verify that the *pMoNIA1* promoter is functional by testing induction with varying nitrate concentrations (5–10 g/L KNO_3_).

### Problem 4

Inconsistent or Variable Phenotypes Across Transformants (related to Expected Outcomes and Step 17)

### Potential solution

- Screen a larger pool of initial transformants (10–15) to identify those with consistent nitrogen-dependent phenotypic switching.
- Perform replicate phenotypic assays on the same transformant line across multiple experiments to distinguish true variability from experimental noise.
- Verify molecular integration by PCR (Figure 3) for all transformants before extensive phenotypic characterization. Only use transformants with confirmed correct integration patterns.
- Critical: For the most rigorous analysis, select 3–5 independent transformants with confirmed correct integration and robust phenotypic switching for use in subsequent experiments. Biological replicates derived from independent transformation events help account for potential positional effects or line-to-line variation.
- Consider that some variability may stem from differences in culture conditions (medium age, temperature fluctuations, light exposure). Standardize all growth parameters when comparing transformants.
- Transformants with intermediate phenotypes may still be useful for certain applications but should be characterized thoroughly and their limitations noted in experimental reports.

### Problem 5

Filter Papers with No Fungal Growth After Co-cultivation (Step 16)

### Potential solution

- Verify that *M. oryzae* spores were viable and at the correct concentration (5.0 × 10^5^ spores/mL). Dead or over-diluted spores will not germinate.
- Ensure co-cultivation medium (IM solid) was prepared correctly with 200 µM acetosyringone and appropriate carbon sources to support both Agrobacterium and fungal growth.
- Check that co-cultivation temperature (25°C) and light conditions (continuous light) were maintained throughout the 48-hour period.
- Avoid over-mixing or crushing spores when spreading them with the bacterial spreader, as mechanical damage can reduce viability.

## RESOURCE AVAILABILITY

### Lead contact

For additional information or requests for resources and reagents, please contact the lead author, Mostafa Rahnama, at.

### Materials availability

The wild-type *M. oryzae* strain and the derived mutants used in this study, as well as the plasmids generated, are available upon reasonable request and may require appropriate permits.

### Data and code availability

This study did not generate any unique datasets or code beyond the experimental results described.

## ACKNOWLEDGMENTS

We thank Dr. Mark Farman (University of Kentucky) for providing the *M. oryzae* strain Guy11 and pMY200 plasmid. We are grateful to Dr. Marc-Henri Lebrun for his expertise in nitrate reductase promoter systems and for guidance in identifying and validating the *M. oryzae* NIA1 promoter (*pMoNIA1*) for conditional gene expression.

This work was supported by funding provided by the Center for the Management, Utilization, and Protection of Water Resources at Tennessee Tech University.

## AUTHOR CONTRIBUTIONS

J.K. executed the experiments. M.R. supervised the study and wrote the manuscript.

